# Sidelobe suppressed Bessel beams for one-photon light-sheet microscopy

**DOI:** 10.1101/2024.08.15.607323

**Authors:** Chetna Taneja, Jerin Geogy George, Stella Corsetti, Philip Wijesinge, Graham D. Bruce, Maarten F. Zwart, Shanti Bhattacharya, Kishan Dholakia

**Affiliations:** SUPA School of Physics and Astronomy and Centre for Biophotonics, University of St Andrews, North Haugh, St Andrews, Fife, KY16 9SS, UK; Department of Electrical Engineering, IIT Madras, Chennai, India; School of Psychology and Neuroscience, Centre for Biophotonics, University of St Andrews, St. Andrews, UK; Centre of Light for Life and School of Biological Sciences, University of Adelaide 5005, Australia

## Abstract

Bessel beams (BB) have found widespread adoption in various forms of light-sheet microscopy. However, for one-photon fluorescence, the transverse profile of the beam poses challenges due to the detrimental effect of the sidelobes. Here, we mitigate this issue by using a computer generated phase element for generating a sidelobe suppressed Bessel beam (SSBB). We then progress to perform a comparison of biological imaging using SSBB to standard BB in a light-sheet geometry. The SSBB peak intensity is more than an order of magnitude higher than the first sidelobe. In contrast to a standard BB light-sheet, SSBB does not need deconvolution and propagates to depths exceeding 400*μ*m in phantom samples maintaining a transverse size of 5 *μ*m. Finally, we demonstrate the advantage of using a SSBB light-sheet for biological applications by imaging fixed early-stage zebrafish larvae. In comparison to the standard BB, we observe a two-fold increase in contrast-to-noise ratio (CNR) when imaging the labelled cellular eye structures and the notochords. Our results provide an effective approach to generating and using SSBB light-sheets to enhance contrast for one-photon light-sheet microscopy.

## I. INTRODUCTION

Light-sheet fluorescence microscopy (LSFM) has proven to be an excellent tool for volumetric imaging of a variety of biological samples [1]. This includes developing embryos, multicellular specimens, and cleared mice brains and is due to its optical sectioning capability and low photo-toxicity while still preserving high resolution[2–6]. LSFM’s distinct advantages arise from its geometry which utilizes a sheet of light for illuminating the sample, with the fluorescence signal collected in a perpendicular direction to the illumination axis[7].

The axial resolution and depth of focus (DOF) of a LSFM system are the two important variables for consideration [8]. The axial resolution determines optical sectioning ability, whereas the DOF defines the imaging volume of the sample [7]. Both these parameters are related to the thickness and beam shape of the light-sheet[8, 9]. The Gaussian beam shape is the most popular choice for creating a light-sheet but its rapid divergence, which is inversely related to its beam waist, limits the DOF we can achieve at any given resolution and provides a major limitation for LSFM [7]. As a result, illumination using ‘propagation-invariant’ beams, particularly the Bessel beam (BB), serves as an alternative due to these fields possessing invariant transverse profiles over extended propagation distances[10]. This leads to a larger DOF whilst maintaining a high axial resolution [11–13].

Propagation invariant BBs have a symmetric sidelobe structure in their transverse profile that has to be considered in the light-sheet imaging process due to their potential to contribute unwanted out-of-plane fluorescence which results in reduced image contrast for one-photon excitation [14, 15]. This contribution of the sidelobes can be suppressed in the case of multi-photon LSFM, due to the nonlinear nature of the imaging process (for example, the quadratic dependence of fluorescence emission on excitation intensity in two-photon fluorescence)[16–18]. However, one-photon light-sheet has advantages due to potential simplicity in use, inexpensive laser choice for more compact arrangements and a wider range of compatible fluorophores [19, 20]. Therefore, an approach is needed to mitigate the effect of sidelobes in one-photon BB LSFM.

In order to improve the signal-to-noise ratio (SNR) in one-photon mode with BB, various post-processing methodologies have been used, including deconvolution[21], content-aware compressed sensing (CACS)[22], deep learning[23], and photo-bleaching imprinting[20]. These methods may be computationally very expensive and complex [24, 25].

An alternative route is to consider modifying the transverse structure of the BB itself to reduce the amplitude of the sidelobes, yet retaining the inherent features of such a beam, namely propagation invariance and self-healing [26–29]. This is termed sidelobe suppressed Bessel beam (SSBB). Several theoretical methods, such as interfering two structured light-sheets[30], superposition of two BBs [31] and self-learning sidelobe elimination [32] have been proposed to generate SSBB light-sheet specifically for LSFM.

Recently, Di Domenico et al. reported sidelobe reduction in BB by projecting a double-ring mask on a spatial light modulator (SLM) (in amplitude mode) to generate SSBB light-sheet [33]. However, the projected ring pattern causes the loss of many incident photons, significantly reducing the incident power. In contrast, Wang et al. used a phase modulation approach for efficient generation of high-contrast BB light-sheet[34]. However, there is an absence of major experimental studies using SSBB for biological imaging in LSFM. In this study, we generate both the BB and the SSBB by encoding an appropriate phase element onto a SLM in a light-sheet setup and present a comparison of the performance of a SSBB LSFM to one using a standard BB. We demonstrate that sidelobe suppression in the SSBB significantly reduces out-of-plane fluorescence generation. Further, we quantify the point spread function (PSF) for our LSFM system. PSF provides a good indication of the image quality for any LSFM system. We observe a reduction of sidelobes in the PSF for SSBB light-sheet without needing deconvolution as in the case of a standard BB light-sheet. This suppression is valid even at depth (>400 *μ*m) in scattering samples making the SSBB ideal for biological imaging. Finally, to demonstrate this relevance, we image fixed and labelled zebrafish larvae (4-5 dpf) using both approaches. Our results show a two-fold improvement in contrast to noise ratio (CNR) using the SSBB light-sheet over its standard BB counterpart. We demonstrate that a SSBB can generate superior image quality to that of the BB, including the BB combined with deconvolution. Our results suggest that the SSBB can enable facile LSFM imaging without additional post processing for routine biological imaging.

## II. MATERIALS AND METHODS

### A. Phase element calculation for SSBB generation

The superposition of two zeroth-order BBs with slightly different wave-vectors (*k*-vector) can be given as

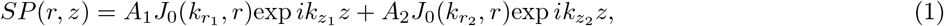

where *J*_0_ is the Bessel function of zeroth-order, and 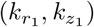 and 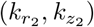 are radial and longitudinal wavevectors of the two different superposing BB. For reduction of sidelobes, the superposed beam’s intensity is suppressed over a transverse region (*r*_0_-*r*_1_) along the propagation distance (*z*_0_-*z*_1_). That is achieved by optimising beam parameters to minimise the following integral:

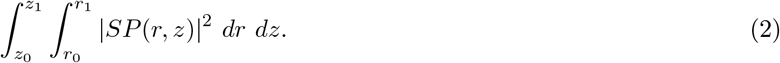

The optimised parameters 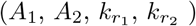 have been utilised to design axion phases for BB/SSBB generation. It is important to note that for the resultant superposition beam, there are two phase terms (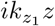 and 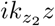) in Eq. 2 which change and coincide repeatedly with distance *z*. This results in oscillatory behaviour of the propagation invariant region with period *P*

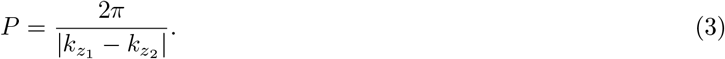

The above equation shows that the period which defines the region of suppressed intensity in the sidelobes can be increased with smaller difference in the two BB’s wavevectors. This results in the superposition beam closely resembling BB with inefficient lobes suppression. It is also evidence that, for SSBB, minimising the integral in Eq. 3 is crucial. This results in an overall reduction of intensity for SSBB in comparison to BB. For constant input power, the generated BB carries more power in comparison to an equal radius SSBB. To take this into account, the SSBB has been used with greater incident power and comparative studies have been performed after normalisation with respect to peak intensities of respective images. A detailed discussion on the SSBB generation method can be found in our recent publication[31].

### B. Image acquisition and processing

The imaging plane (*x, y, z*) is positioned at 45° angle with respect to the sample plane (orthogonal coordinates *x*^′^, *y*^′^, *z*^′^). The sample is translated along *x*^′^ axis capturing a series of images. To transform the co-ordinates from imaging plane to physical coordinates, we utilise the affine transformation function in ImageJ. For an image stack represented by a matrix M with pixel coordinates (*n, m, l*) with camera pixel size *δp* and translation distance *δx*^′^, the transformation into the physical (sample) plane is given by: *x*^′^ = *n δp* + *l*sin 45^*°*^, *y*^′^ = *m δp, z*^′^ = *l δp* sin 45^*°*^.

### C. Sample preparation and staining

#### 1. Phantoms

14 *μ*l volume of 1 *μ*m diameter green fluorescent microspheres (fluoro-max, G0100) with particle density 1.05 gm/*cm*^3^ was used without any dilution and mixed with 1.5% agarose solution. To prepare Phantom, 60 *μ*l volume of 1 *μ*m diameter non-fluorescent microspheres (Duke Standards, LOT No.- 247589) with the same particle density is added to the same agarose solution. The samples were pipetted and placed in a single cell of an 8-cell sample mount for imaging.

#### 2. Zebrafishes

Zebrafish larvae expressing GCaMP6 (4-dpf) were euthanised using MS222 (250 mg/L, Sigma Aldrich E10521, ethyl 3-aminobenzoate methanesulfonate) and fixed overnight in 4% paraformaldehyde (PFA) at 4^*°*^C and subsequently washed 3 times in phosphate buffered saline (PBS).

## III. RESULTS

### A. Experimental set-up for BB and SSBB LSFM

Figure 1(a) shows the simplified schematic of the optical setup used for the measurements. A 488 nm laser beam passes through a half-wave plate (HWP) to align input polarisation with the direction of liquid crystals for maximising the SLM efficiency. Simulated phase masks for the generation of both BB/SSBB are projected onto the SLM. The BB is generated by projecting an axicon phase with an opening-angle, *α*_1_= 0.36°. For the SSBB, the approach for phase mask generation has been described earlier[31]. Briefly, the SSBB is generated by a combination of two axicon phase functions (*α*_2_= 0.26° and *α*_3_ = 0.46°) into a single phase function using a random multiplexing technique [31]. These two phase functions correspond to two BBs with slightly different wave-vectors (*k*-vectors) which are superposed such that the beam’s transverse intensity is suppressed throughout the defined propagation distance. Both beams have been generated with equal core radius of *R*= 60 *μ*m just after the SLM. The depth of focus (DOF) for the BB depends on the input Gaussian beam radius and opening-angle of the axicon. For the BB, the DOF after the SLM is *∼*175 mm. SSBB is generated periodically with a period *P* [26, 31], where *P* determines the propagation invariant distance over which the sidelobes are suppressed below the desired range (see methods). Ideally, a large distance in which the sidelobes are suppressed is desired (similar to the BB’s DOF). The period can be increased by decreasing the difference between the wave-vectors of individual BB, but at the expense of the efficiency of lobes-suppression. In our case, the optimised value for *P* for an efficient sidelobes suppression (*<* 5% of peak intensity in first sidelobe) has been calculated to be around 90 mm.

**FIG. 1.**
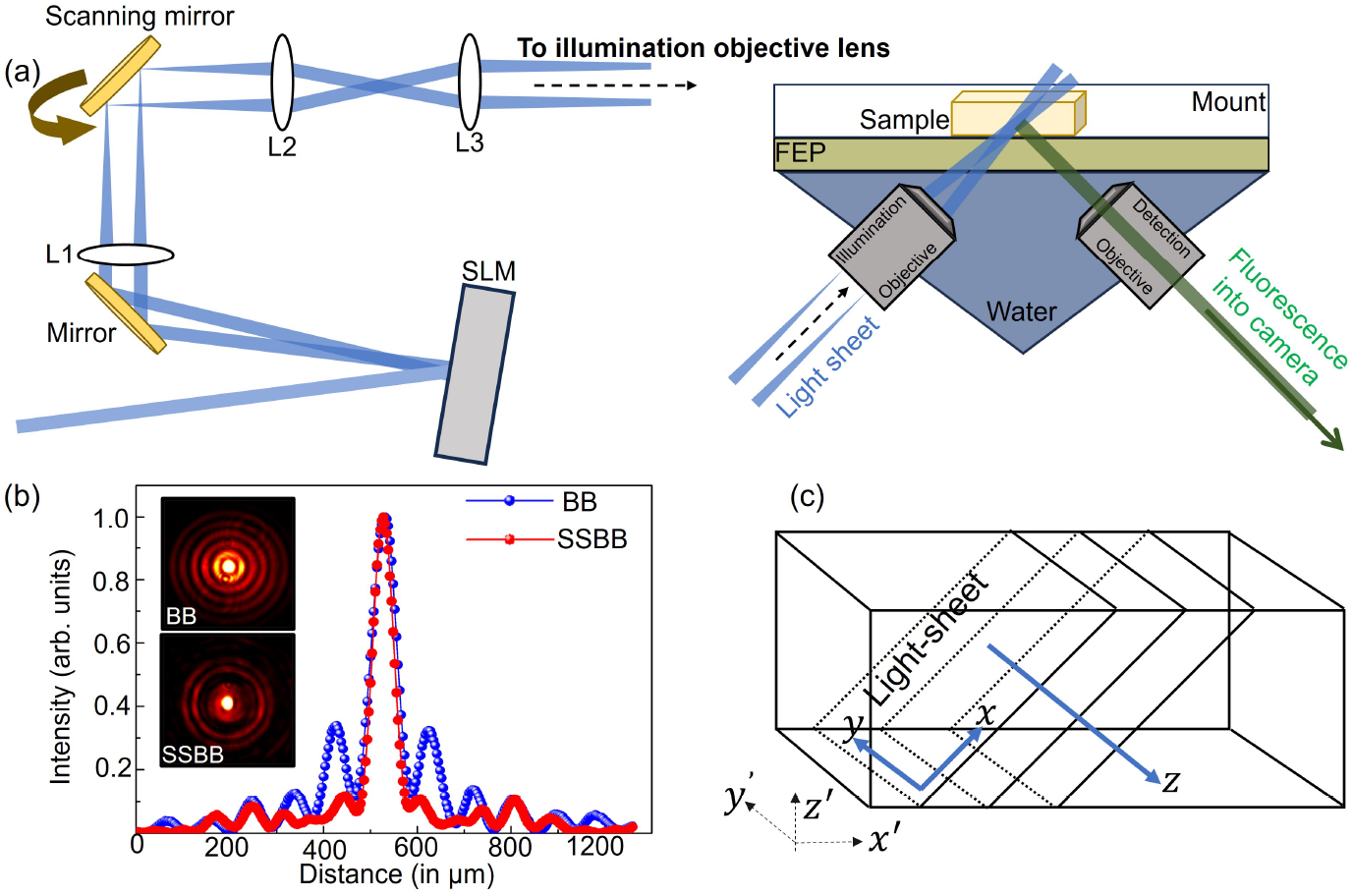
(a) Light-sheet fluorescence microscopy experimental setup. Laser: 488 nm (Toptica iBeam Smart, 100 mW), L1-3: lenses (*f* = 100, 50, 75 mm), SLM: Spatial light modulator (LCOS, Hamamatsu), illumination objective: 0.367NA 4X (Navitar). Inset: schematic of the sample positioning. The specimen is placed inside an 8-well mount lined with fluorinated ethylene propylene (FEP) film (RI= 1.33) to match the refractive index of the water filling the chamber. Detection objective: 0.367NA 4X (Navitar), Camera: Iris15 (Photometrics). (b) Intensity profile along the transverse cross-section of the beam for the BB (blue curve) and the SSBB (red curve). Inset shows images of the BB and the SSBB generated just after the SLM. (c) Definition of two sets of coordinates in the LSFM system; with respect to the light-sheet acquisition (*x,y,z*) and the physical sample plane (*x*^′^,*y*^′^,*z*^′^).

Figure 1(b) shows the transverse intensity line-profiles measured after the SLM for the generated BB (blue curve) and SSBB (red curve). Intensity profiles for both the beams are shown as the insets in Figure 1(b). For the BB, the first two sidelobes contribute to 19% and 8% of the peak intensity respectively, whereas the SSBB carries less than 5% of the peak intensity in the first two sidelobes.

For the LSFM set-up, a lens (L1) is used to project the Fourier transform of the BB (single ring) and the SSBB (two concentric rings) onto a galvanometer mirror to create a scanning light-sheet (along *y*-axis). Lenses L3 and L4 are used to project the scanned light-sheet to the back focal plane of the water-immersion objective lens. Inset shows the positioning of the sample. The specimen is mounted using an 8-well mount with fluorinated ethylene propylene (FEP) film placed inside to match the refractive index of the water filling the chamber hosting the illumination and detection objectives. The detection objective, which has the same specifications as the illumination objective, collects the signal emitted by the sample. This signal is focused onto the camera using a tube lens (TL) after being filtered using a notch filter (*λ*_*c*_= 488 nm) and a band-pass filter (*λ*_*c*_= 520 nm, Δ*λ*= 60 nm).

The light-sheet propagates along the *x*-axis with an amplitude set by the scanning mirror in the *y*-axis. The detection axis is along *z*-axis (Figure 1(c)). Another set of coordinates (*x*^′^, *y*^′^, *z*^′^) defines the system with respect to the imaged sample. The sample mount is placed on a translation stage and moved along *x*^′^-axis with a high-resolution motorised linear actuator (PI M-230.10) to collect an image stack. To transform the image stack collected from light-sheet to physical (sample) coordinates, we use affine transformation function in the software tool ImageJ.

### B. BB and SSBB light-sheet characterization in fluorescein

Firstly, we image the BB and SSBB in fluorescein solution. The *xy* cross-section of both the BB and the SSBB intensity profiles obtained by imaging their propagation in a fluorescein solution (concentration ≈ 30 *μ*M) are shown in Figure 2(a). For comparison, the images are individually normalised with respect to their own profiles. For the BB case, the sidelobes around the intensity maxima at *y*= 0 *μ*m generate a significant amount of out-of-plane fluorescence in both *y* < 0 and *y* > 0 directions. Such fluorescence is suppressed in the SSBB case.

**FIG. 2.**
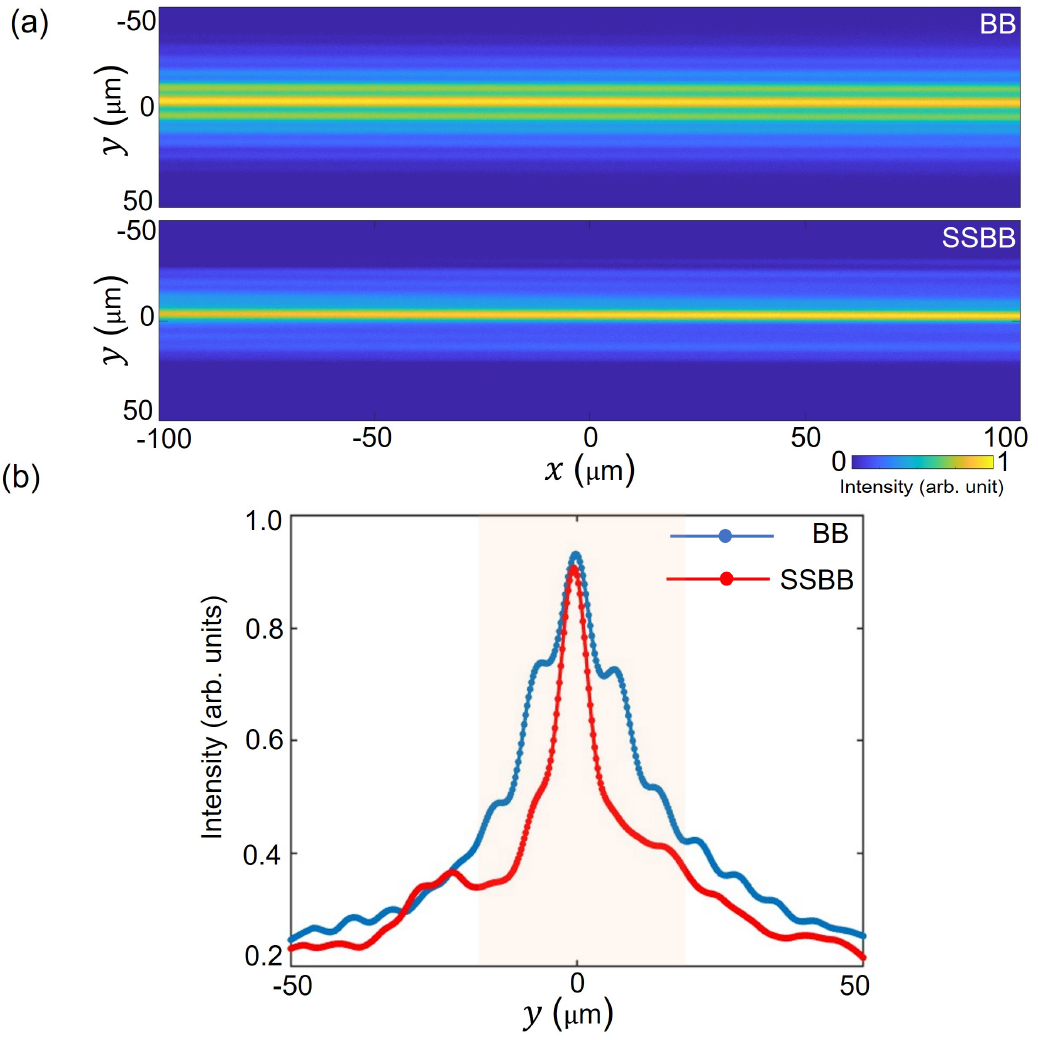
(a) *xy* sections of the BB and SSBB intensity profiles in fluorescein solution. Both images are normalised with respect to their maximum intensity. (b) Intensity profiles for the BB (blue curve) and the SSBB (red curve) obtained by plotting the line profiles at *x* = 0 *μ*m along the *y*-axis.

For a further qualitative analysis, the BB and the SSBB intensity profiles at *x*= 0 *μ*m are compared in Figure 2(b). The fluorescence intensity peaks generated by the sidelobes around the central intensity maximum for the BB are not visible for the SSBB. The fluorescence intensity peak by the first sidelobe of the BB is only 25% lower than that of the central-core maxima, which in the SSBB light-sheet has been measured to be reduced by more than 50% along with the reduction of the fluorescence contribution from all other lobes.

Fluorescein images are used to experimentally characterize the BB and the SSBB light-sheet. The ‘waist’ of the light-sheet at the sample plane *r* is calculated from the thickness of the central-lobe of beam’s transverse profile in Figure 2(b). For both the BB and SSBB, thickness of the light-sheet is found to be *r*= 4.80 ± 0.32 *μ*m from five independent measurements.

The telescopes in the excitation path reduce the DOF of the BB at the sample plane to *∼*1.4 mm which matches with the field-of-view (FOV= 1.5 mm) of the camera along the detection pathway. As described above, SSBB is generated periodically with a DOF at the sample plane of around 0.54 mm. Nonetheless, it is important to note that the SSBB still offers almost two-fold increase in DOF when compared to a Gaussian beam with the same core radius.

### C. Lobe suppression in the PSF of the SSBB LSFM

To demonstrate the capability of the SSBB light-sheet for imaging, we compare the PSF of the microscope using both the BB and the SSBB light-sheets. To determine the PSF, 400 nm diameter green-fluorescent beads embedded in agarose are imaged and analyzed. This size was chosen to be below the diffraction limit of the system. Image stacks (100 images) are collected by translating the sample along the *x*^′^-direction with a step-size of 0.5 *μ*m. The exposure time is set to 500 ms for each step. Selected *xy* cross-sectional image of the beads collected using both the BB and SSBB light-sheets are shown in Figure 3(a) and (b) respectively. Insets (i) and (ii) show the lateral (*xy*) and axial (*xz*) profiles of a single bead marked with the white rectangular box in Figure 3(a) and (b). The comparison between the normalised intensity (along the black dashed line in the inset) for the lateral profiles (Figure 3(c)) show overlapping curves for the BB (Blue) and the SSBB (red). This provides similar values for the full-width half maximum (FWHM). The value of the FWHM determines the lateral resolution of the system (along *x, y*-axis). We analysed 10 beads for each of the BB and SSBB light-sheet and obtained the same lateral resolution of *∼*1.33 ± 0.05 *μ*m for each beam profile, as expected.

**FIG. 3.**
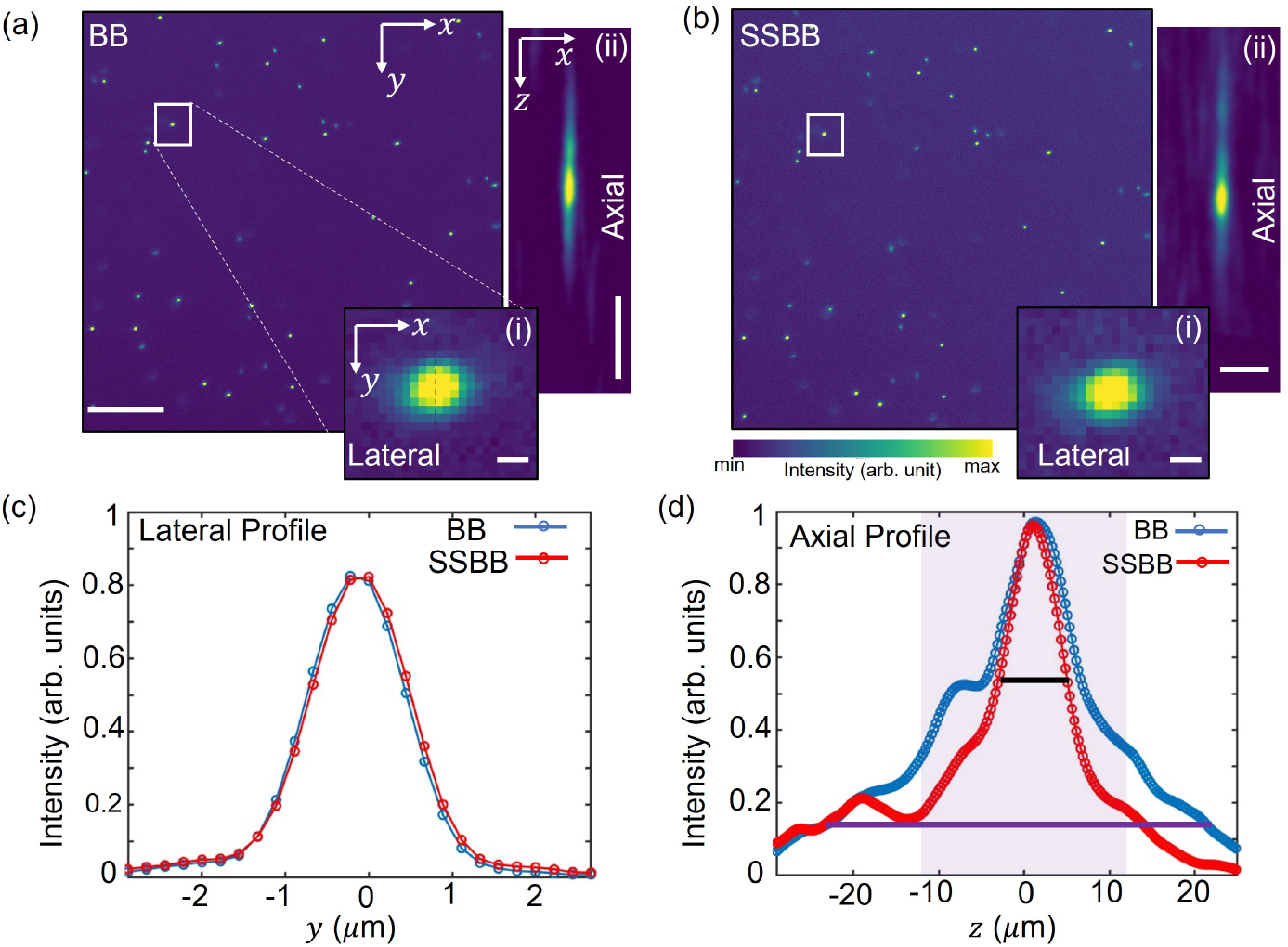
Representative *xy* cross-sectional images of 400-nm diameter fluorescent microspheres embedded in agarose acquired with the (a) BB and (b) SSBB light-sheet. Scale bar: 50 *μ*m. Insets (i) in Figure 5(a) and (b) show a magnified view of beads marked by the white box. Scale bar: 1 *μ*m. Insets (ii) in Figure 5(a) and (b) show the *xz* view of the same bead. Scale bar (along *z*- and *x*-axis): 10 *μ*m. (c-d) Plot of the line profiles through the beads in the insets (i) and (ii). The curves for the BB and SSBB are represented in blue and red, respectively. The solid black and purple lines in (d) show the width of the curve at the FWHM and 1*/e*^2^ value, respectively.

For the axial resolution, the width of the bead profile along the detection axis is measured for both the BB and the SSBB. The intensity profile along the *z*-axis of the bead (inset a-b) for the BB (blue) and the SSBB (red) are shown in Figure 3(d). For the BB, the sidelobes in the PSF (along *z*-axis) can be seen. The distance between the central maxima and first sidelobe (7 *μ*m) matches with the distance calculated in the BB light-sheet profile in fluorescein (see Figure 2 (b)). For the SSBB, these side-lobes are suppressed and thus do not contribute to the cross-sectional image of the bead. The FWHM of these curves provides the axial resolution of the LSM system. At the FWHM, the width of each curve is very similar, with only 16% reduction for the SSBB (solid black line). This results in a similar axial resolution for both beams (∼ 6.92 ± 0.15 *μ*m for SSBB, ∼ 6.99 ± 0.16 *μ*m for BB). An important thing to note is the overall reduction of the PSF using the SSBB. The width of the PSF decreases by more than 45% in comparison to the BB at 1/*e*^2^ of the maximum intensity (solid purple line). This suggests that for LSM, the SSBB light-sheet would provide significantly higher contrast.

### D. Lobe suppression in the PSF with the BB with deconvolution and SSBB light-sheet

The axial profiles of the imaged beads show the utility of the SSBB light-sheet to suppress the sidelobes in the PSF of LSFM system (Figure 3(d)). For a traditional BB light-sheet in one-photon mode, image post-processing tools such as deconvolution are utilised for suppression of these sidelobes. Next, we compare the PSF of the bead imaged with a standard BB light-sheet to the PSF calculated with employing deconvolution (orange curve) and using a SSBB light-sheet (red curve) (Figure 4a). Deconvolution of the BB was performed using the Richardson-Lucy method implemented by DeconvolutionLab2[35].

**FIG. 4.**
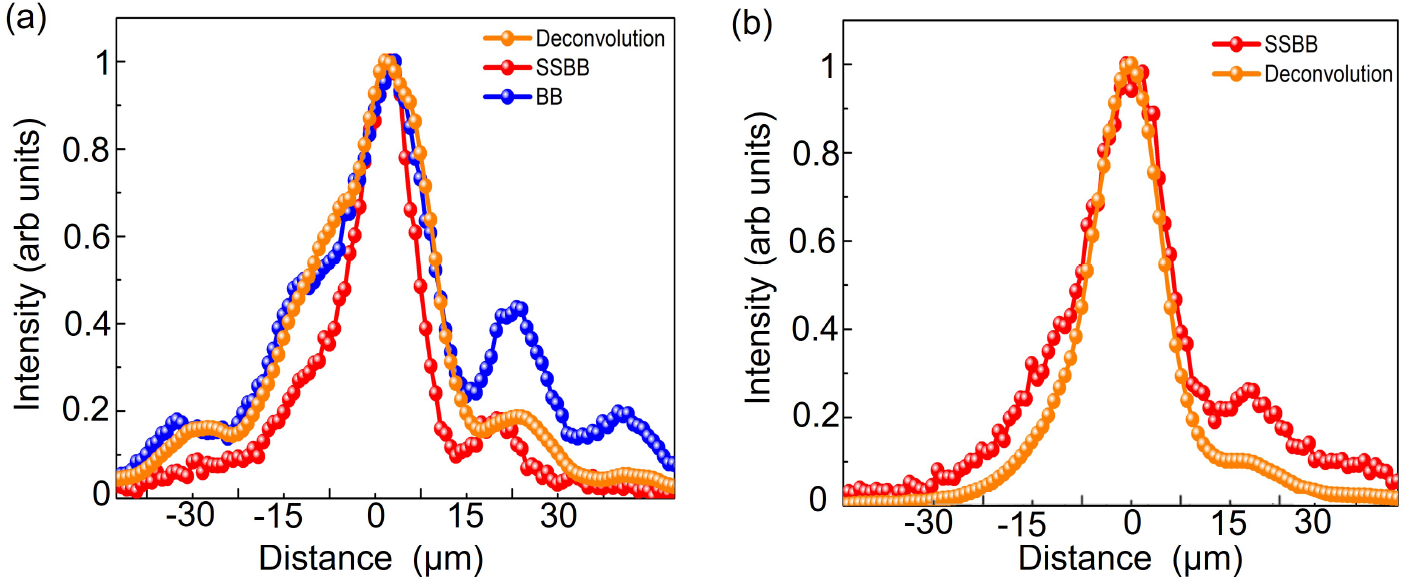
(a) Intensity profile along the *z*-axis of a 400 nm diameter green fluorescent bead imaged with BB (blue curve), BB with deconvolution (orange curve) and SSBB (red curve) (b) Comparison between the intensity profile of a single bead imaged with SSBB (red curve) and SSBB with deconvolution (orange curve).

The PSF calculated with deconvolution provides a smoother curve in comparison to the BB light-sheet, however could not effectively suppress the sidelobes. This is similar to the previous findings on using deconvolution on propagation invariant beam shapes [21]. On the other hand, SSBB provides better suppression of sidelobes and overall reduction of the PSF width without needing any additional analysis and complex computational methods.

Additionally, we check whether the performance of the SSBB light-sheet could further be improved by utilising deconvolution. Figure 4(b) shows the line-profiles of the PSF imaged with SSBB light-sheet with (orange curve) and without deconvolution (red curve). The effect of the sidelobes are further suppressed with utilising deconvolution, however SSBB itself provides effective lobe suppression for the LSFM system.

### E. Imaging phantoms: sustaining the transverse profile of the SSBB at depth

As the light-sheet penetrates deeper into a sample, scattering plays an increasingly important role in the quality of the recorded signal. This is a major challenge for imaging at depth and brings into question whether the transverse profile for both the BB and the SSBB are sustained in the imaging process. Therefore, to compare the performance of the BB and the SSBB light-sheet at depth in a scattering medium, we imaged phantoms (mimicking a scattering biological sample) prepared with a high concentration of non-fluorescent (scattering) beads mixed with a low concentration of green fluorescent beads (each of diameter of 1 *μ*m), embedded in agarose (approx size 1 mm^3^). Details are given in the methods section. Figure 5(a) shows images of a single *xy* plane of phantom from an image stack acquired with both the BB and the SSBB light-sheets. In both cases, the fluorescence intensity attenuates along the *x*-axis due to scattering from non-fluorescent beads. To quantify the sidelobe suppression at depth, we recorded the intensity projection along the *xz* plane for the BB and the SSBB (Figure 5(b)). The axial profiles (line profile along *z*-axis) of beads at different depths within the sample imaged with both light-sheets (BB- blue, SSBB- red curve) are compared in Figure 5(c). It is clear that the SNR, defined as the ratio of intensity maxima to background intensity, for both the beams decreases with depth.

**FIG. 5.**
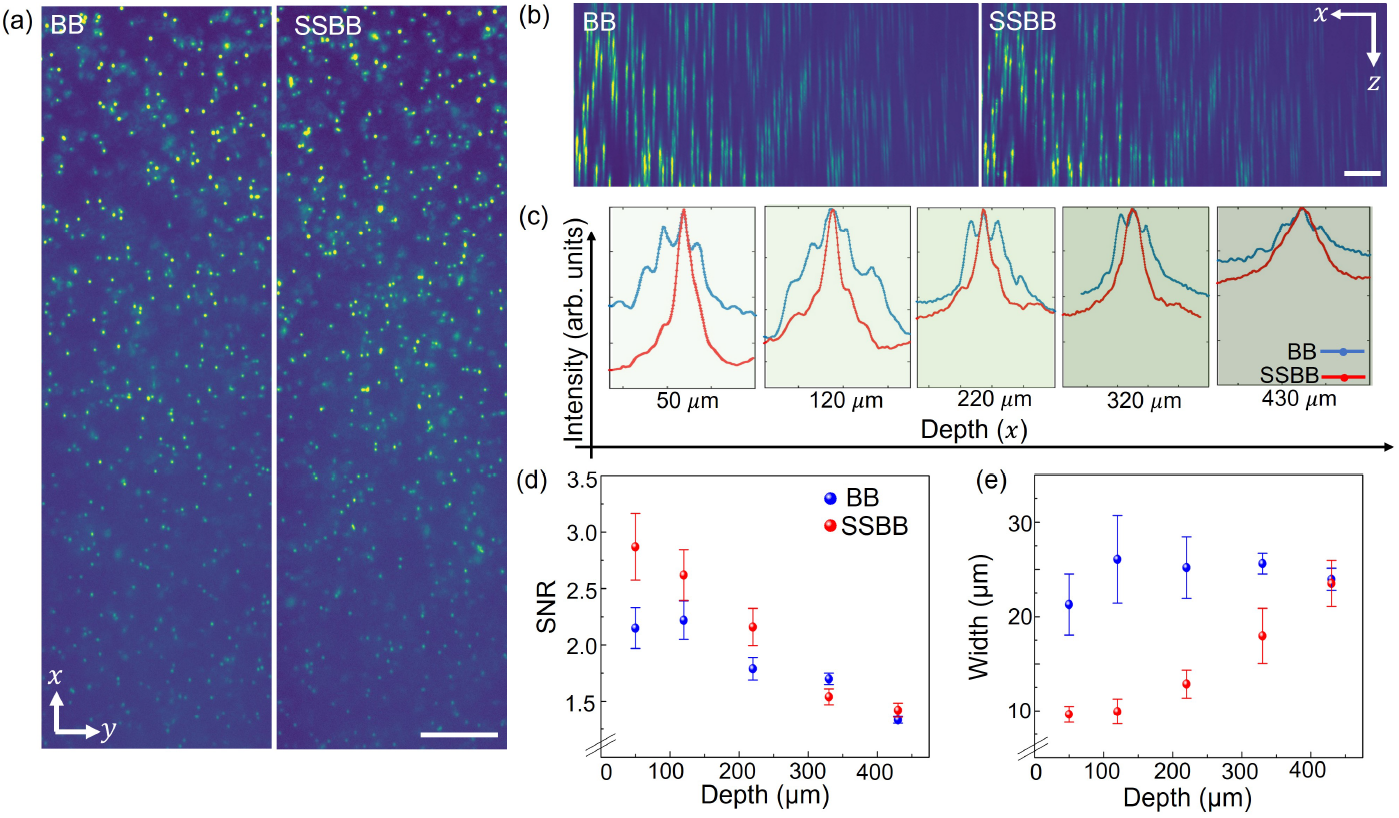
(a) *xy* cross-sectional plane image of a mixture of 1 *μ*m diameter green-fluorescent and 1 *μ*m diameter non-fluorescent beads embedded in agarose imaged using the BB and the SSBB. Scale bar: 50 *μ*m. (b) Intensity projections in the *xz* plane imaged with the BB and SSBB light-sheets. Scale bar: 50 *μ*m. (c) Comparison between the axial profiles of random beads imaged using the BB (blue curve) and the SSBB (red curve) as a function of depth. Going deeper into the sample increases the background (scattering) from the non-fluorescent beads for both BB/SSBB, however lobes suppression in PSF is still visible at 430 *μ*m in depth. (d) Signal-to-noise (SNR) for the imaged beads as a function of depth. (e) Width of the line profiles of the imaged beads in phantom as a function of depth.

We compare the SNR from analysing the axial profiles of 10 beads at each depth value imaged using both the BB and SSBB light-sheet (Figure 5(d)). The standard deviation is shown as the error bars for each data point. It can be noted that SSBB provides a better SNR in comparison to BB light-sheet for imaging closer to the FEP film. However, at greater depths (>220 *μ*m) both light-sheets give similar values for SNR. For the SSBB, SNR ranges from a value of 2.87 ± 0.6 at *x*= 50 *μ*m near the FEP surface, to 1.42 ± 0.2 at *x*= 430 *μ*m depth. This can be attributed to the presence of scattering in the sample (due to non-fluorescent beads) which attenuates the fluorescent signal significantly and the scattering contributes to increased noise.

Another important point to note is the width of the curves shown in Figure 5(e) for different depth values. It can be seen that the width of the bead profile increases with depth. It is challenging to calculate the axial resolution at a function of depth from these axial profiles due to increased scattering which results in sidelobes contributing to the FWHM. For a traditional BB, the contribution of the sidelobes is more prominent than the SSBB light-sheet which results in a greater value of the width at half maximum for the intensity profiles of the beads. Although SSBB provides reduced width for imaging closer to the surface, it increases to almost being equal to BB light-sheet at greater depth values. This deterioration of the value of the width is expected as SSBB is more sensitive to the scattering than BB light-sheet due to being generated by the superposition of two BB.

However, it is important to note the lobe suppression for SSBB light-sheet, which is visible even at depths up to 430*μ*m with SSBB. This proves that for samples where scattering plays a crucial role such as biological specimens, SSBB would provide better contrast images.

### F. Enhanced contrast in zebrafish imaging

Finally, to assess contrast improvement when imaging biological samples, we imaged zebrafish larvae (4-5 dpf) labelled with GCaMP using both the BB and the SSBB light-sheet modalities. Zebrafish is ideal for this application. Their body is transparent during larval development, which allows them to be imaged at depth [36]. Further, zebrafish are a common and relevant model organism for studying diseases and understanding larvae development processes [37, 38]. In this context, light-sheet imaging of zebrafish with improved image quality would be very valuable for medical research [39]. For our present studies, structures labeled with GCaMP, having peak excitation around 480 nm, are imaged with fluorescence emission at 510 nm.

We first labelled fluorescent cellular structures around the lens of the zebrafish eye. Images for this eye can be seen in the *xy* cross-sectional plane images acquired with the BB (Figure 6(a)) and the SSBB (Figure 6(b)) light-sheet approaches. For comparison, both images are individually normalised with respect to the peak intensity of the individual image. The blue and red boxes depict the outer layers of an eye, which is closest to the surface of the FEP film. The insets (c) and (d) provide an enlarged view of the same region. When imaging with SSBB, there is an improvement in contrast compared to the BB, with both outer layers (outer nuclear layer (ONL) and photoreceptors) being easily distinguishable with less background. Along with the double layer, the cellular structures (retinal ganglion cell layer (RGL), inner plexiform (IPL) and inner nulclear layer (INL)) of the eye are also much more visible.

**FIG. 6.**
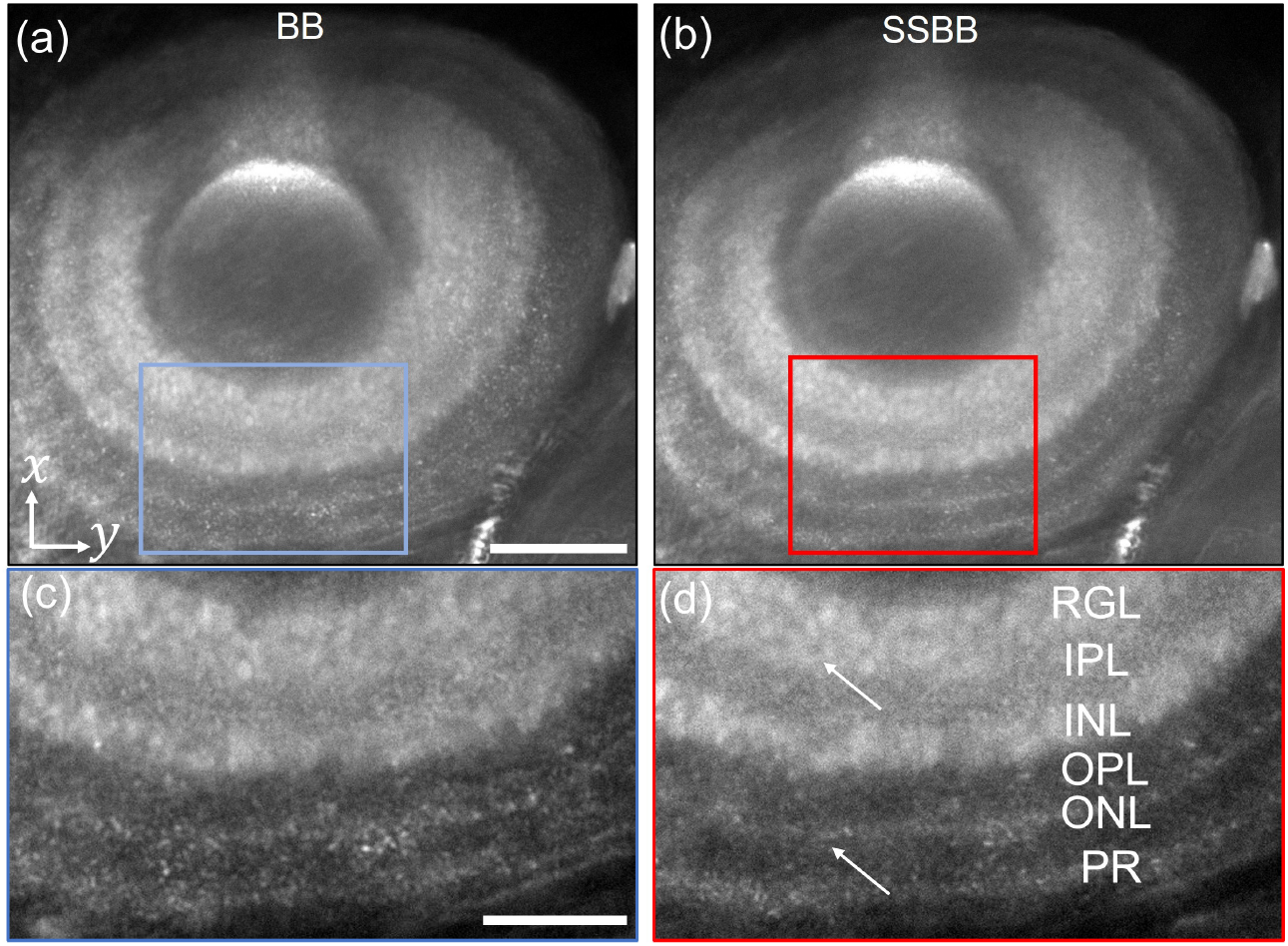
Imaging GCaMP labelled cell nuclei of the zebrafish (4-5 dpf) eye using (a) BB and (b) SSBB. Scale bar: 50 *μ*m. Insets (c) and (d) show magnified view of rectangular box in (a) and (b), respectively. Scale bar: 25 *μ*m. RGL, retinal ganglion cell layer; IPL, inner plexiform layer; INL, inner nuclear layer; OPL, outer plexiform layer; ONL, outer nuclear layer; PR, photoreceptors. White arrows show imaged regions with improved contrast using the SSBB light-sheet.

For both images in the insets we calculated the contrast-to-noise ratio, which is defined as 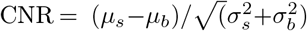. Here, *μ*_*s*_ and *μ*_*b*_ denote signal and background mean intensities, respectively, while *σ*_*s*_ and *σ*_*b*_ represent the standard deviations of signal and background intensity. This metric for quantifying contrast or image quality for biological samples has been widely used in the literature [40].

The image acquired using the BB light-sheet yields a CNR of ∼1.53, while the image acquired using the SSBB light-sheet gives a higher CNR of ∼3.02. Other areas, in addition to those displayed in the insets, also exhibit distinct background noise suppression, resulting in an overall improvement in image quality when employing the SSBB.

A similar increase in CNR is observed when imaging different regions of the zebrafish (notochord and muscles) with the SSBB light-sheet. The *xy* cross-sectional images (normalised and presented with the same gray scale) acquired with the BB and SSBB light-sheets are shown in Figure 7(a) and (b), respectively. The increase in contrast is visible when different structures, such as the notochord and muscle, can be easily identified from the SSBB light-sheet image. Figure 7(c) shows the line profiles along the black dashed line depicted in Figure 7(a) for both the images. For BB (blue curve), there are multiple peaks with no clear distinction of the structures. In the case of SSBB (red curve), the intensity peaks are clearly distinguishable and have smaller widths in comparison to BB. Furthermore, for SSBB the overall background decreases significantly, resulting in a much clearer image.

**FIG. 7.**
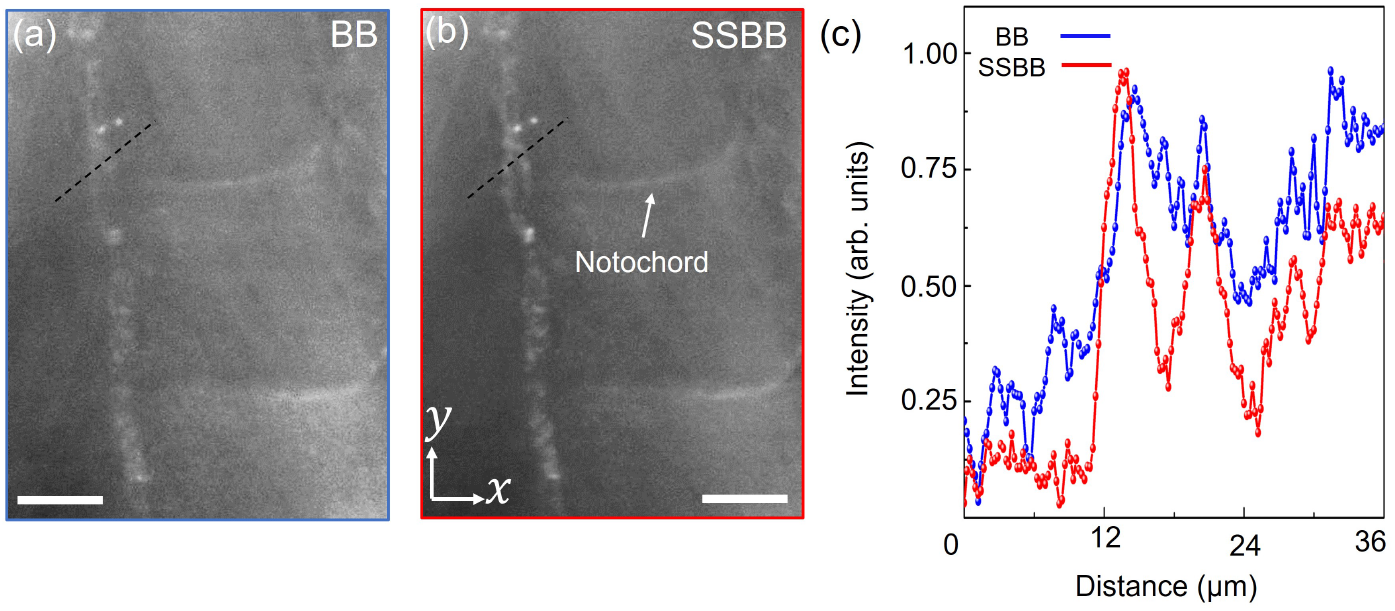
Imaging the region around notochord of a fixed and labelled zebrafish using (a) BB and (b) SSBB light-sheets. Scale bar: 25 *μ*m. (c) Line-profile along the black dotted line shown in (a) for BB (blue curve) and SSBB (red curve).

## IV. DISCUSSION AND CONCLUSION

Imaging of one-photon fluorescence with the SSBB for LSFM will be very useful for advancing optical imaging for biomedical applications. Here, we answered the unexplored question of ‘how good is the SSBB in comparison to a standard BB for biological imaging in the LSFM?’. We utilise an efficient generation method of using a phase mask which can also easily be adaptable to integration with meta-optical elements [31]. The generated SSBB achieves more efficient suppression of the sidelobes in the PSF of the LSFM system, even when compared to a deconvolution post-processed BB light-sheet. Additionally, our results provide a comparative study for the BB and the SSBB LSFM in biological imaging. We show a two-fold contrast improvement for the SSBB from imaging cellular structure of an eye for fixed zebrafish larvae (4-5 dpf) labelled with GCaMP with further improved contrast demonstrated in notochord and muscle images.

Previously published results show the averaged two-fold reduction in the background by imaging a mixture of different diameter fluorescent spheres embedded in agarose without implementation of SSBB light-sheet for biological imaging [33]. We use CNR as a metric to quantify the image quality since CNR provides contrast between tissue of interest relative to the background scattering (surrounding tissues) and readily used as a powerful tool for biological image analysis. We also show that generated SSBB sustains its transverse profile up to 430 *μ*m deep into the sample which is another important requirement for high-resolution volumetric imaging for biological samples.

One important feature of this generation method is the periodicity associated with the SSBB. Specifically for LSFM, this periodicity could prove to be advantageous for high-resolution imaging of larger biological specimen through simultaneous imaging different regions from multiple SSBB light-sheets. The optical set-up could be modified easily (changing the focal length of a lens in the illumination path) to generate multiple SSBB with reduced DOF within the FOV of the imaging camera. Reduced DOF also ensures better suppression of the sidelobes and hence provides better axial resolution. Other configurations are also possible for utilising the SSBB’s periodicity such as using multiple cameras for different SSBB light-sheets. These techniques could be highly beneficial for obtaining a high-resolution 3D reconstruction of larger biological samples using LSFM.

## ACKNOWLEDGMENTS

We thank Dr. Pierce Mullen, Eleonora Gagliardi from the School of Psychology and Neuroscience, University of St Andrews for providing labelled zebrafish larvae samples.

## DATA AVAILABILITY

The research data underpinning this publication will be accessed at https://doi.org/10.17630/92a0783e-2fd3-43ca-bce7-ccfeb0bd5732

